# Differential Impact of Doxorubicin Dose on Cell Death and Autophagy Pathways during Acute Cardiotoxicity

**DOI:** 10.1101/2022.01.15.476450

**Authors:** Philip Kawalec, Matthew D. Martens, Jared T. Field, Wajihah Mughal, Andrei Miguel Caymo, Donald Chapman, Bo Xiang, Saeid Ghavami, Vernon W. Dolinsky, Joseph W. Gordon

## Abstract

Doxorubicin (DOX) is an effective anthracycline used in chemotherapeutic regimens for a variety of haematological and solid tumors. However, its utility remains limited by its well-described, but poorly understood cardiotoxicity. Despite numerous studies describing various forms of regulated cell death and their involvement in DOX-mediated cardiotoxicity, the predominate form of cell death remains unclear. Part of this inconsistency lies in a lack of standardization of *in vivo* and *in vitro* model design. To this end, the objective of this study was to characterize acute low- and high-dose DOX exposure on cardiac structure and function in C57BL/6N mice, and evaluate regulated cell death pathways and autophagy both *in vivo* and in cardiomyocyte culture models. Acute low-dose DOX had little impact on cardiac structure or function; however, acute high-dose DOX elicited substantial cardiac necrosis resulting in diminished cardiac mass and volume, with a corresponding reduced cardiac output, and without impacting ejection fraction or fibrosis. Low-dose DOX consistently activated caspase-signaling with evidence of mitochondrial permeability transition. However, acute high-dose DOX had only modest impact on common necrotic signaling pathways, but instead led to an inhibition in autophagic flux. Intriguingly, when autophagy was inhibited in cultured cardiomyoblasts, DOX-induced necrosis was enhanced. Collectively, these observations implicate inhibition of autophagy flux as an important component of the acute necrotic response to DOX, but also suggest that acute high-dose DOX exposure does not recapitulate the disease phenotype observed in human cardiotoxicity.

## Introduction

Doxorubicin (DOX) is an effective chemotherapeutic agent used in many haematological and solid tumors (1, 2). DOX functions primarily as a DNA intercalator, thereby obstructing biosynthesis of new macromolecules, and secondarily as a topoisomerase II inhibitor, leading to disruption of topoisomerase II-mediated DNA repair and eventually cell death (3). This ability to damage DNA and disrupt cellular growth has made DOX an attractive chemotherapeutic agent (4). DOX serves as an essential pillar of chemotherapeutical treatment of sarcomas, carcinomas, and haematological neoplasms, being one of the most actively used agents in the treatment of breast cancer, ovarian cancer, and lymphoma (5).

Despite its widespread use and effectiveness as an anti-neoplastic agent, DOX efficacy remains complicated in the clinical environment due to its well-described, but poorly understood cardiotoxic effects. DOX-mediated cardiotoxicity manifests as arrhythmias, systolic and diastolic dysfunction, dilated cardiomyopathy, and ultimately congestive heart failure, which has a mortality rate of approximately 50% (6, 7). The incidence of heart failure amongst patients receiving either DOX or DOX-derivatives has been reported to be as high as 26%, and can occur acutely during treatment or a number of years after finishing chemotherapy (8). Many of the investigations into the mechanisms of DOX-induced cardiotoxicity are based on research studies utilizing various cardiomyocyte cell- and animal-models. Much of the focus has been on elucidating the cardiomyocyte response to DOX and the pathogenesis of DOX-induced cardiomyopathy to better direct prevention and treatments. Many pathways have been implicated in this process, including oxidative/REDOX stress, mitochondrial dysfunction, autophagy/mitophagy, dysfunctional iron metabolism, regulated cell death, and extracellular matrix remodelling (9–13). (11, 14).

Currently, an emphasis has been placed in determining which of the many regulated cell death processes are responsible for cardiomyocyte death after DOX exposure (15). Whereas apoptosis was once thought to be the major form of regulated cell death in DOX-mediated cardiotoxicity, novel regulated cell death mechanisms such as mitochondrial permeability transition-dependent necrosis, necroptosis, pyroptosis, and ferroptosis, have been shown to have important, if not predominant roles in cell death induced by DOX (15–18). To date, there is no consensus as to which form of regulated cell death is dominant, but it is highly probable that several are induced at the same time and are likely context dependent. In addition, autophagy is the process by which cells destroy redundant or defective cellular products, and has become a major point of interest in the area of DOX-mediated cardiotoxicity. Despite a large number of studies published on the subject of whether autophagy is protective or detrimental to cardiomyocytes when exposed to DOX, the matter is still unclear (16).

At least some of the complexity in our understanding of DOX-induced cardiotoxicity and regulated cell death can be attributed to the lack of standardized DOX dosages, duration of DOX treatment, and choice of models used for *in vivo* and *in vitro* experiments, which is further complicated with respect to both species and biological sex as most DOX studies have been exclusively performed in male rodents. In general, three dosing regiments have been used in animal models to study DOX-induced cardiotoxicity, including acute or chronic designs where animals are treated with: 1) one high dose of DOX acutely (19), or 2) one high-dose of DOX and surviving animals are evaluated longitudinally (20), and 3) multiple lower doses of DOX over the span of several days to weeks (21, 22). Although, it is worth noting that the 5-day mortality following a single high-dose DOX treatment (ie. 20mg/kg) is roughly 50-60% (19, 20), which seems disproportionally excessive compared to human studies on acute DOX-induced cardiotoxicity (8). Currently, studies use doses ranging from 5-25 mg/kg, with the lower dosages used in chronic exposure and higher dosages being in an acute dose setting (22–25). Moreover, it has also been suggested that the cumulative life-time DOX exposure is one of the most predictive variables in determining DOX-induced cardiotoxicity, which suggests that acute high-dose DOX exposure is comparable with chronic low-dose DOX exposure (26, 27). Regarding model choice, a recent review by Christidi & Brunhum (2021) found that studies investigating DOX cardiotoxicity typically use a combination of neonatal rat cardiomyocytes, adult rat cardiomyocytes, Sprague Dawley rats, C57BL/6J -N mice, or H9c2s cardiomyoblasts, where a few use a single model, and only one used all five (16, 22), which raises concerns regarding whether observations are translatable amongst the different dosing routines and models.

This non-standardized approach has resulted in discordant, and at times contradictory, findings in the literature. The objective of this study was to characterize how acute low- and high-dose DOX treatments impact mouse cardiac structure and function, as well as evaluate recently identified cell death pathways in several cardiomyocyte models, including C57BL/6N mice, primary neonatal rat cardiomyocytes and H9c2 rat cardiomyoblasts, with the aim to more completely understand the acute adverse effects of DOX cardiotoxicity.

In this report, we describe the effects of acute low-dose DOX (LDD; 8mg/kg) and high-dose DOX (HDD; 20mg/kg) on cardiac structure/function, cell death and autophagy pathways. Acute HDD elicited substantial cardiac necrosis determined by serum troponin levels, while echocardiography revealed a diminished cardiac output and mass, without impacting ejection fraction or fractional shortening. Evaluation of cell death pathways identified that acute LDD activated caspase signalling, while acute HDD inhibited autophagic flux and increased BNIP3 expression - which was not observed in a chronic DOX model. When autophagy was inhibited in cultured cardiomyoblasts, DOX-induced necrosis was enhanced. In addition, DOX exposure elicited a dose-dependent increase in mitochondrial ROS and lipid peroxidation in culture, and inhibition of ferroptosis with ferrostatin-1 provided a modest increase in cell viability. Collectively, these observations help characterize the structure/function and gene expression alterations in the heart during the acute phase of DOX-induced injury. Our findings also suggest that acute HDD does not recapitulate the disease phenotype observed in human cardiotoxicity and that not all changes in gene expression are conserved between HDD and chronic DOX exposure models.

## Materials and Methods

### In Vivo High and Low Dose DOX Mouse Models

All procedure involving animal studies were approved by the Animal Care Committee of the University of Manitoba, which diligently follows the principles of biochemical research involving animals put in place by the Canadian Council on Animal Care. C57BL/6N mice were bred and housed in our facility, and randomly allocated to one of three groups at 10 weeks of age: control (saline, n=3), low-dose DOX (LDD; 8mg/kg, n=3), or high-dose DOX (HDD; 20mg/kg, n=4). For the chronic DOX model, eight-week-old C57BL/6N mice received weekly intraperitoneal injections of 8 mg/kg/week of DOX, or weekly intraperitoneal injections of equal volume 0.9% saline, for a total of 4 weeks, as described previously (21). Injections with the appropriate saline or DOX dose were delivered intraperitoneally and mice were weighed and inspected daily. On post-injection day (PID) 3, echocardiography was performed on all mice using a Vevo 2100 High-Resolution Imaging System equipped with a 30-MHz transducer (RMV-716; Fujifilm-VisualSonics, Toronto ON) to assess cardiac morphometry and function in B-mode, M-mode and Doppler imaging, as described previously (28). During imaging, mice were placed on a heated ECG platform to maintain body temperature at 37C and were sedated under mild anesthesia (induction with 3% isofluorane and 1 L/min oxygen and maintained at 1-1.5% isofluorane and 1 L/min oxygen). Data was collected and interpreted by a trained echocardiographer and measurements averaged over four cardiac cycles. Mice were sacrificed and serum and cardiac tissues were collected for western blot analysis and histological analysis. Serum Troponin-I was measured using the Ultra-Sensitive Mouse Cardiac Troponin-I ELISA Kit purchased from Life Diagnostics (CTNI-1-US; Life Diagnostics; West Chester, PA).

### In Vitro Cell Culture

H9c2 cells (ATCC CRL-1446) were cultured in Dulbecco’s modified Eagle’s medium (DMEM; Hyclone) media with 10% Fetal bovine serum (Hyclone), and penicillin/streptomycin, as described previously (28–32). Cells were incubated at 37 °C with 5% CO_2_. Primary ventricular neonatal cardiomyocytes (PVNCs) were isolated from 2-3 day old Sprague Dawley rats using a Pierce Primary Cardiomyocyte Isolation Kit from ThermoFischer (#88281) and cultured according to the manufacturer’s instructions (28, 30–32). Both H9c2 cells and PVNCs were treated with either vehicle control, 1μM, 5μM, or 10μM of DOX (Sigma Aldrich, reconstituted in sterile double distilled water), which was added directly into the culture media.

### Live Cell Imaging

H9c2 or PVNCs were cultured and treated with DOX for 24 hours, after which they were incubated with the appropriate dye for 30 minutes, washed with PBS three times, and then imaged in culture media. All imaging was done using a Zeiss Axiovert 200 inverted microscope equipped with a Calibri 7 LED Light Source (Zeiss) and Axiocam 702 mono camera (Zeiss). Calcein-AM, ethidium homodimer-1, tetramethylrhodamine methyl ester (TMRM), and Hoechst 33342 dyes were purchased from Biotium. MitoSOX was purchased from Life technologies while LysoTrackerRed DND-99, Rhod-2AM, and dichlorodihydrofluorescein diacetate (H_2_-DCFDA; DCF) were purchased from Invitrogen. Mitochondrial permeability transition pore imaging (mPTP) imaging was carried out with Calcein-AM and cobalt chloride using the same method previously described by our lab (28, 30, 31). Mito-pHred, a plasmid-based biosensor (33) and dihydro-Rhod2 (28) imaging were described previously.

### Immunofluorescence

Following DOX treatment, H9c2 or PVNC cells were washed and fixed with 4% paraformaldehyde for 15 minutes at 37°C. Cells were washed with PBS three times for 5 min and incubated with blocking buffer consisting of 5% goat serum (CST) and 0.3% Triton X-100 (Sigma Aldrich) in PBS. Cells were incubated overnight with the appropriate antibody in 1% bovine serum albumin (BSA; Sigma) and 0.3% Triton X in PBS. The next day cells were washed with PBS three times for 5 min and incubated with the appropriate secondary antibody for one hour. Cells were washed three more times with PBS, mounted onto slides using ProLong™ Diamond Antifade Mountant with DAPI (Invitrogen), and imaged using the Axiovert 200 microscope (28).

### Western Blotting

Cell lysates were collected, analysed for protein content, and prepared for electrophoresis in the same manner as described previously (33). Samples were resolved on acrylamide gels and then transferred to PVDF membranes overnight. Membranes were then blocked with 5% powdered milk or BSA in Tris-buffered saline with Tween-20 (TBST) for 1 hour at room temperature before being incubated with the appropriate antibody in 1% powered milk or BSA in TBST. The membranes were then washed with TBST three times for 15 minutes and incubated with the appropriate secondary antibody in 1% powdered milk or BSA in TBST for 90 minutes. The blots were then washed three more times with TBST for 15 minutes and protein was visualized using SignalFire Plus ECL (CST) and a BioRad Chemidoc.

### Antibodies

The following antibodies were used for western blotting and immunofluorescence: HMGB1 (CST #3935), Rodent-specific Bnip3 (CST #3769), Actin (sc-1616), BNIP3L/Nix (CST #12396), p53 (CST #9282), Tom20 (CST #42406), SQSTM1/p62 (CST #23214), LC3B (CST #83506), BCL2L13 (proteintech 16612-1-AP), Parkin (CST #2132), AIF XP (CST #5318), Phospho-RIP3 (CST #91702), Cleaved Caspase-3 (CST #9664), Cleaved-PARP (CST #67495), Cleaved Gasdermin D (CST #10137), GPX4 (CST #52455), normal rabbit IgG (sc-2027), normal mouse IgG (sc-2025), Alexa-Fluor 647 anti-mouse (Jackson 715-605-150), Alexa-Fluor 647 anti-rabbit (Jackson 711-605-152), Alexa-Fluor 488 anti-rabbit (Jackson 711-545-152).

### Real-time PCR

Total RNA was extracted from pulverized frozen tissue or from cultured cells by TRIzol method. Following column purification using Qiagen RNeasy kit and DNase treatment, cDNA was generated with QScript cDNA super mix (Quanta BioSciences). RT-PCR was performed using PerfeCTa SYBR green super mix on a ABI 7500 Real-Time PCR Instrument, and normalized to β-actin expression, as described previously (30).

### Transfections and Plasmids

H9c2s were cultured and transfected with plasmids using JetPRIME Reagent (PolyPlus). After incubation with the transfection media for 6 hours, the media was replaced with normal culture media and incubated overnight at 37°C. The next day, the cells were treated with DOX and imaged 24 hours later. GW1-Mito-pHRed (Addgene 31474) was a gift from Gary Yellen (34); and pEGFP-LC3 (Addgene, 24920) was a gift from Toren Finkel (35).

### Annexin-PI Flow Cytometry

H9c2 cells were pretreated with various cell death inhibitors: 1μM Ferrostatin-1 (FER1; Sigma 0583), 5nM Bafilomycin A1 (BAF; Sigma 1793), or 50μM zVAD(OMe)-fmk (ZVAD; Enzo BML-P416-0001) in culture media for 4 hours. After the pre-treatment was finished, media was replaced with new media which contained the same inhibitor along with the appropriate concentration of DOX. The cells were incubated for 24 hours, after which they were collected and stained according to the BD Pharmingen FITC Annexin V Apoptosis Detection Kit. Flow cytometry was run on a Attune NxT Acoustic Focusing Cytometer (28, 36).

### Statistical Analysis

Data are presented in graphs as mean ± standard error. Experiments were analyzed using a 1-way ANOVA with subsequent Tukey test for multiple comparisons. All cell culture experiments were performed from 3 independent passages or isolations. All statistical analysis was done using GraphPad Prism software. p<0.05 = *, p<0.01 = **, p<0.001 = ***, p<0.0001 = ****.

## Results

### Acute DOX Exposure Elicits Cardiomyocyte Necrosis and Reduced Cardiac Size

To compare the effects of acute low-dose DOX (LDD; 8 mg/kg) and acute high-dose DOX (HDD; 20 mg/kg) on cardiac structure and function, C57BL/6N mice were treated by intraperitoneal injection (Figure 1A). Control mice received an equivalent volume of saline. Three days post-injection, mice underwent echocardiography and tissue collection. We chose 3-day post injection since our objective was to evaluate cardiac structure and function prior to lethality and the 5-day mortality for mice receiving 20 mg/kg DOX was reported to be 50-60% (19). Mice treated with LDD lost on average less than 1 gram of body mass; however, mice treated with HDD on average lost more than 4 grams, or approximately 20 % of their initial mass, suggesting that HDD is acutely toxic (Figure 1B). Consistent with this, HDD treated mice displayed a reduced cardiac mass by 43% when corrected for tibial length, while cardiac mass in LDD treated mice was reduced by 33% but this failed to reach statistical significance (p=0.14). In addition, we evaluated cardiac necrosis through serum Troponin-I, and observed a dosedependent increase with DOX exposure (Figure 1D). Next, we performed Masson Trichrome staining of cardiac sections. Whole heart mounts confirmed that acute DOX treatment reduced cardiac size, particularly in HDD treated animals (Figure 1E). However, higher magnification revealed only a modest increase in perivascular or interstitial fibrosis in the HDD treated mice (Figure 1E).

**Figure 1.**
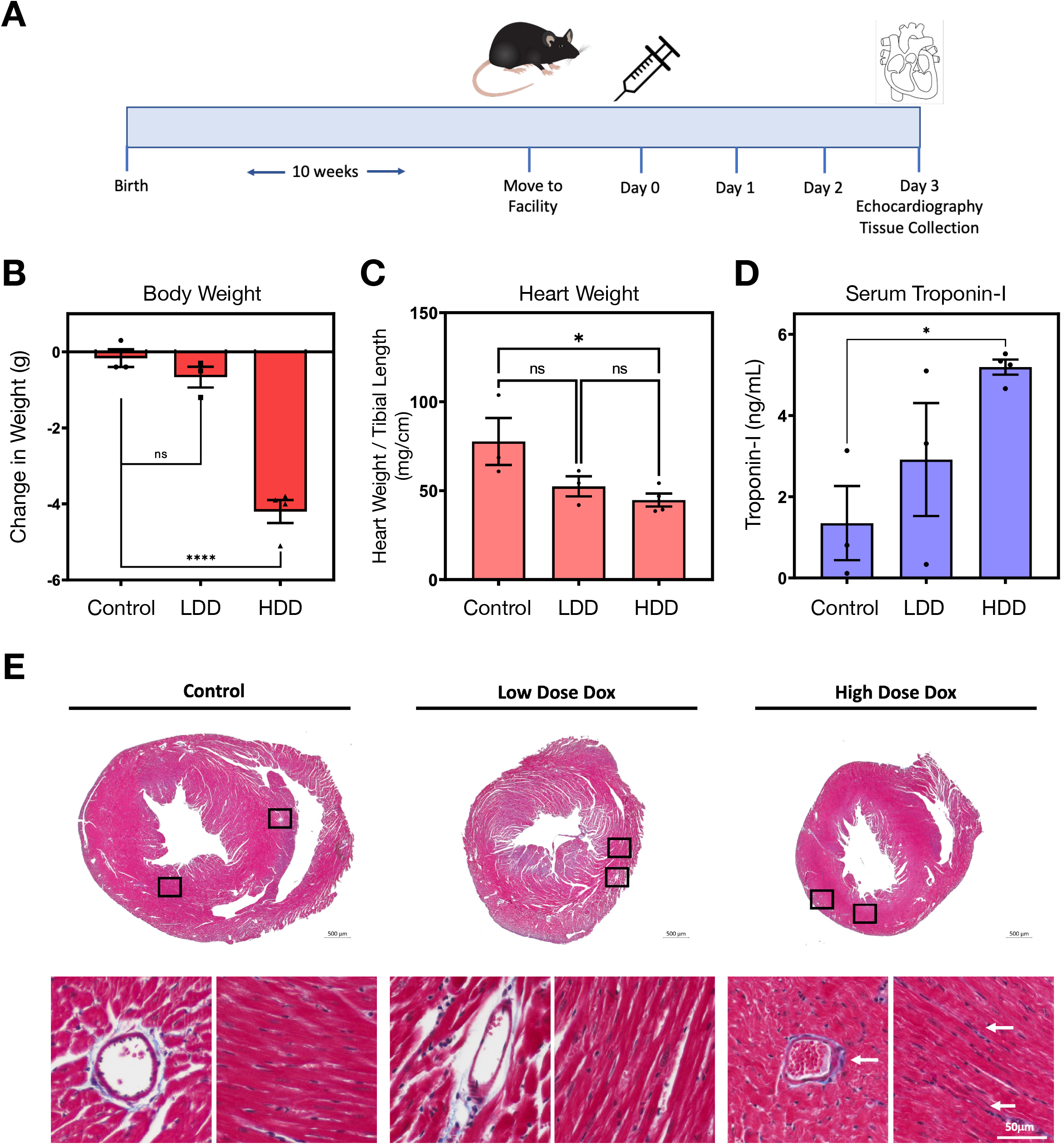
Acute DOX Exposure Elicits Cardiomyocyte Necrosis and Reduced Cardiac Size. A) Schematic of the mouse model of doxorubicin (DOX) exposure, where 10-week old mice received intraperitoneal injections of DOX or saline control. Three days post-injection (PID3), mice are subjected to echocardiography and tissue were harvested. B) Differences in body mass measured in grams (g) from Day 0 until Day 3 (PID3) in control, low-dose DOX (LDD; 8 mg/kg), and high-dose DOX (HDD; 20 mg/kg). C) Differences in heart mass corrected for tibial length (mg/cm) measured at PID3 in control, LDD, and HDD exposed mice. D) Serum Troponin-I evaluated in control, LDD, and HDD. E) Masson Trichrome staining of hearts from Control, LDD, and HDD treated mice. Whole-mounts in transverse sections shown above, and higher magnification perivascular and myocardium/interstitial sections shown below. All data are represented as mean ± S.E.M. * indicates p<0.05, **p<0.01, ***p<0.001, ****p<0.0001. ns = not significant.

### High-dose DOX Treatment Reduces Stroke Volume without Impacting Ejection Fraction

Next, we performed echocardiography on mice 3-days post-DOX injection. We did not detect changes in resting heart rate between the control, LDD and the HDD treated groups (Figure 2A). However, we did observe both a dose-dependent decrease in cardiac output (CO) and stroke volume (SV) in the DOX treated mice (Figure 2B). As an estimate of left ventricle function, we determined ejection fraction and fractional shortening, but did not observe a difference in these parameters in DOX-treated mice, compared to control (Figure 2C). To evaluate if other parameters were contributing to the reduced stroke volume, we evaluated left ventricle volumes and diameters during systole (s) and diastole (d). Shown in Figure 2D and -E, HDD treatment reduced both left ventricle volume and diameter throughout the cardiac cycle, while LDD had no impact on these measurements. We also observed that HDD reduced left ventricle posterior wall thickness (Figure 2F), suggesting that HDD reduced both left ventricle chamber size and wall thickness. We also observed that HDD increased intraventricular relaxation time (IVRT; Figure 2G), suggesting some degree of diastolic dysfunction. Interestingly, we did not observe an impact of either LDD or HDD on the ratio of early to late ventricular filling, determined by the ratio of early peak diastolic annular motion (e’) to late peak diastolic annular motion using tissue Doppler mode (e’/a’; Figure 2H), or pulse wave mode (E/A; not shown), as filling was equally affected in both early and late phases of diastole. In addition, the E/e’ ratio was not significantly different between groups (not shown). Finally, we evaluated the myocardial performance index (MPI), which incorporates parameters of both systolic and diastolic function to provide an index of global ventricular function. Notably HDD significantly increased MPI, which indicates greater cardiac dysfunction (Figure 2I). Collectively, the *in vivo* echocardiography analysis indicates that even though acute HDD presents with heart failure with preserved ejection fraction (HFpEF), as has been suggested to occur in some cases of human DOX-induced cardiotoxicity associated with diastolic dysfunction (37), this model fails to recapitulate the dilated cardiomyopathy observed in human studies (6, 9, 10). In fact, acute HDD results in a smaller heart with reduced ventricular volumes and wall thickness, resulting in reduced stroke volume and slower diastolic filling.

**Figure 2.**
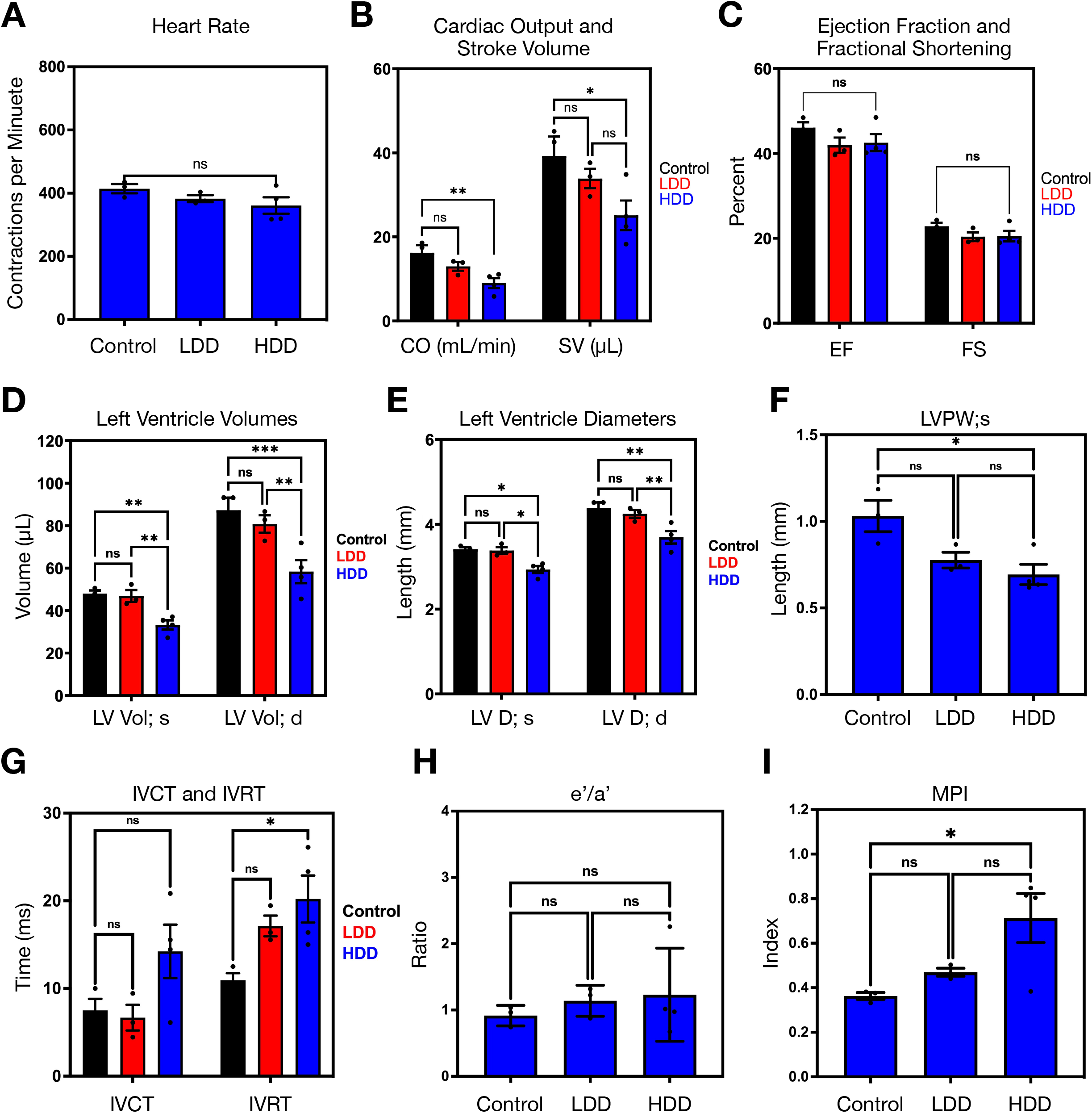
Echocardiography and Doppler Analysis of Low-Dose and High-Dose DOX Treated Mice. A) Heart rate expressed as contractions per minute in control, low-dose DOX (LDD; 8 mg/kg), and high-dose DOX (HDD; 20 mg/kg) treated mice at PID3. B) Cardiac Output (CO; mL/min) and Stroke Volume (SV; μL). C) Calculated Ejection Fraction (EJ) and Fractional Shortening (FS). D) Left ventricular volumes (LV Vol; μL) during systole (s) and diastole (d). E) Left ventricular diameters (LV D; mm) during systole (s) and diastole (d). F) Left ventricle posterior wall (LVPW) thickness during systole (s). G) Intraventricular contraction time (IVCT) and relaxation time (IVRT) measure in milliseconds (ms). H) The ratio of early peak diastolic annular motion (e’) to late peak diastolic annular motion (a’) in tissue Doppler mode. I) Myocardial performance index (MPI) in control, LDD, and HDD treated mice at PID3. All data are represented as mean ± S.E.M. * indicates p<0.05, **p<0.01, ***p<0.001. ns = not significant.

### LDD and HDD Differentially Active Regulated Cell Death Pathways

Using cardiac protein extracts from DOX exposed mice, we next evaluated the activation of regulated cell death pathways, including apoptosis, necroptosis, pyroptosis, and ferroptosis (Figure 3A, -B). We observed an increase in cleaved-caspase 3 expression in the LDD treated hearts, which was not elevated beyond control levels in the HDD treated mice (Figure 3A, -B). Cleaved-caspase 3 is an effector caspase involved in the apoptotic cascade; however, it can also be activated during mitochondrial permeability transition-regulated necrosis concurrent with outer mitochondrial membrane rupture. We did not observe an increase in p-RIP3 in either LDD or HDD treated mice, suggesting that necroptosis is not activated at this time point in DOX-induced cardiotoxicity (Figure 3A, -B). In addition, we observed a modest increase and modest decrease in cleaved-gasdermin D and GPx4 in HDD treated mice, respectively, suggesting that pyroptosis and ferroptosis were activated (Figure 3A, -B). However, these changes in protein expression seemed small in comparison to the magnitude of the increase in serum troponin-I in the HDD group. Finally, we observed a marked increase in the death gene BNIP3 in the HDD group, without an increase in the homologous BNIP3L (ie. Nix) (Figure 3A, -B). Using primary ventricular neonatal cardiomyocytes (PVNCs) in culture, we confirmed some of these expression changes (Figure 3C). In PVNCs, we observed that cleaved-caspase 3 expression was increased in cells treated for 24-hours with 1μM and 5 μM DOX, but not in cells treated with 10 μM DOX. As an indicator of caspase activity, we also observed a concurrent increase in the down-stream target cleaved-PARP1. In addition, we observed a decrease in GPx4 and SIRT3 expression in cells treated with 10 μM DOX, but not in the lower doses (Figure 3C). SIRT3 expression has been previously shown to be decreased in rodent models of chronic DOX exposure (21), and our observation that it is also decreased in acute high-dose exposure, appears to be consistent with the notion that the cumulative exposure to DOX is an important factor predicting cardiotoxicity. Thus, we tested if BNIP3 expression was also increased in a chronic DOX model. Shown in Figure 3D and -E, 4-weeks of 8 mg/kg/week DOX exposure had no impact on BNIP3 expression. Furthermore, we confirmed this by assessing BNIP3 mRNA by realtime PCR (Figure 3F). These observations suggest that not all changes in gene expression are consistent between acute HDD and cumulative chronic DOX exposure.

**Figure 3.**
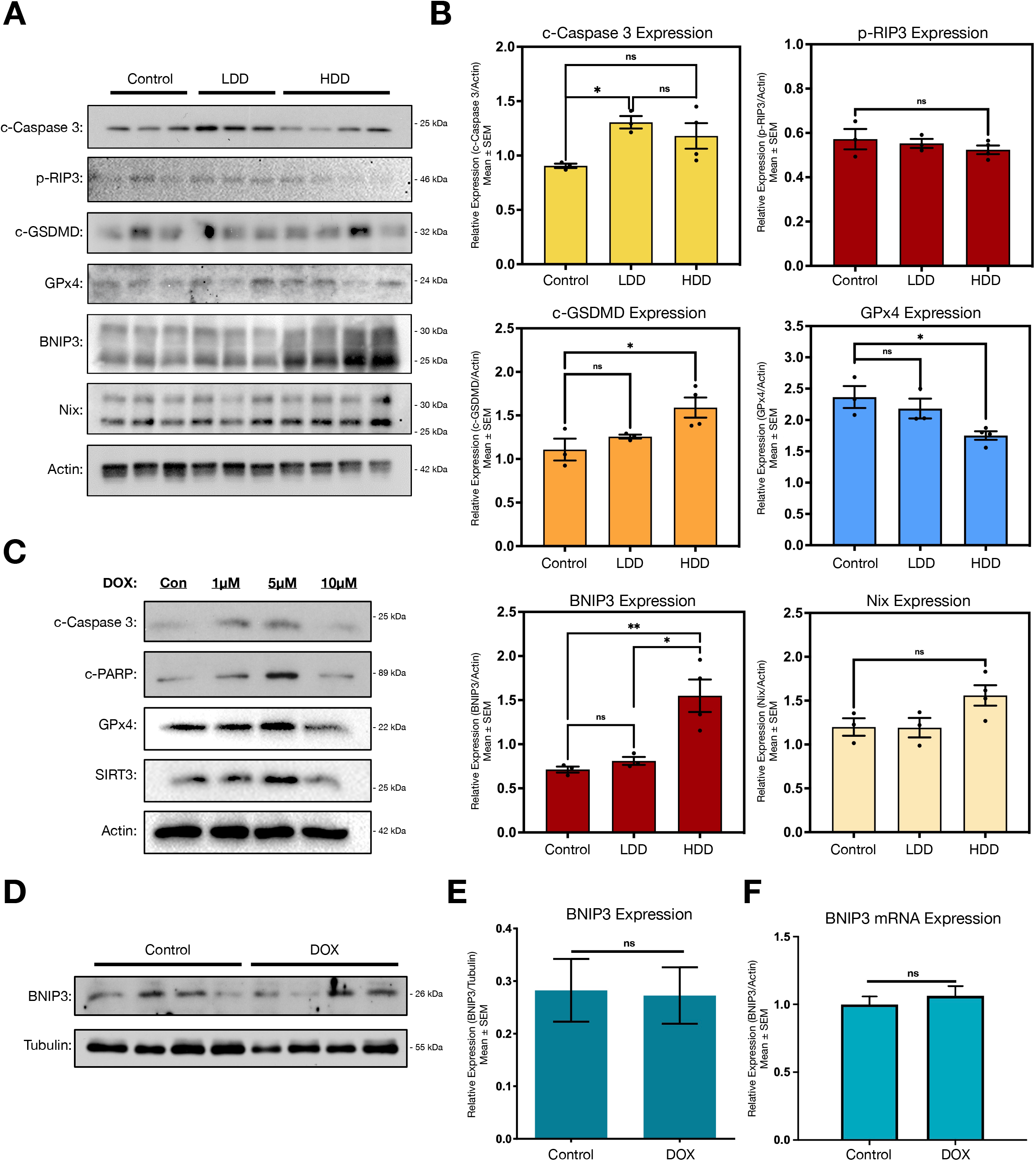
Differential Activation of Regulated Cell Death Pathways by LDD and HDD. A) Western blot analysis of control, low-dose DOX (LDD; 8 mg/kg), and high-dose DOX (HDD; 20 mg/kg) treated mice at PID3, including cleaved caspase 3 (c-caspase 3), phopho-RIP3 (p-RIP3), cleaved gasdermin D (c-GSDMD), Glutathione Peroxidase 4 (GPx4), and β-actin (Actin). B) Quantification of western blots in (a) relative to β-actin (Actin). C) Western blot analysis from primary ventricular neonatal cardiomyocytes treated with 1, 5, and 10 μM DOX for 24-hours, including cleaved PARP1 (c-PARP). D) Western blots of cardiac extracts from mice treated with DOX for 4-weeks of 8 mg/kg/week or vehicle control. E) Quantification of (d) relative to β-tubulin (tubulin). F) Real-time PCR of cardiac extracts from samples in (d). All data are represented as mean ± S.E.M. * indicates p<0.05, **p<0.01. ns = not significant.

### DOX Dosages Differentially Impacts Mitochondrial Permeability Transition Despite Elevations in Mitochondrial Calcium

Next, we used PVNCs and the H9c2 cardiomyoblast cell line to assess cell death mechanisms in response to increasing DOX exposure. In both H9c2 cell and PVNCs, we observed a dose-dependent decrease in cell viability in live/dead assays, using calcein-AM to stain live cells and ethidium homodimer-I to stain cells with a permeabilized plasma membrane, which typically occurs during necrotic cell death (Figure 4A-C). Interestingly, H9c2 cells were more sensitive to DOX toxicity than PVNCs (Figure 4A-C). As accumulation of mitochondrial calcium has been implicated as an important regulator of cardiomyocyte necrosis, we used reduced Rhod2-AM (ie. dihydroRhod2) to stain calcium within the mitochondria. In both H9c2 cell and PVNCs we observed an increase in mitochondrial calcium with the greatest staining in cells treated with 10 μM DOX (Figure 4D, -E). To confirm calcium accumulation in the mitochondria, we used the transfectable matrix-targeted calcium biosensor, mito-CAR-GECO, in H9c2 cells, and observed increased mitochondrial calcium in response to DOX, especially at higher concentrations (Figure 4F). Using the calcein-AM/cobalt chloride method, we evaluated mitochondrial permeability transition in response to DOX exposure in H9c2 cells and PVNCs. Intriguingly, we observed that 1 μM DOX reduced mitochondrial staining, suggesting permeability transition; however, mitochondrial puncta remained at higher concentrations of DOX despite higher mitochondrial calcium staining (Figure 4G-I). These observations suggest that alternate necrotic mechanisms may operate at high concentrations of DOX.

**Figure 4.**
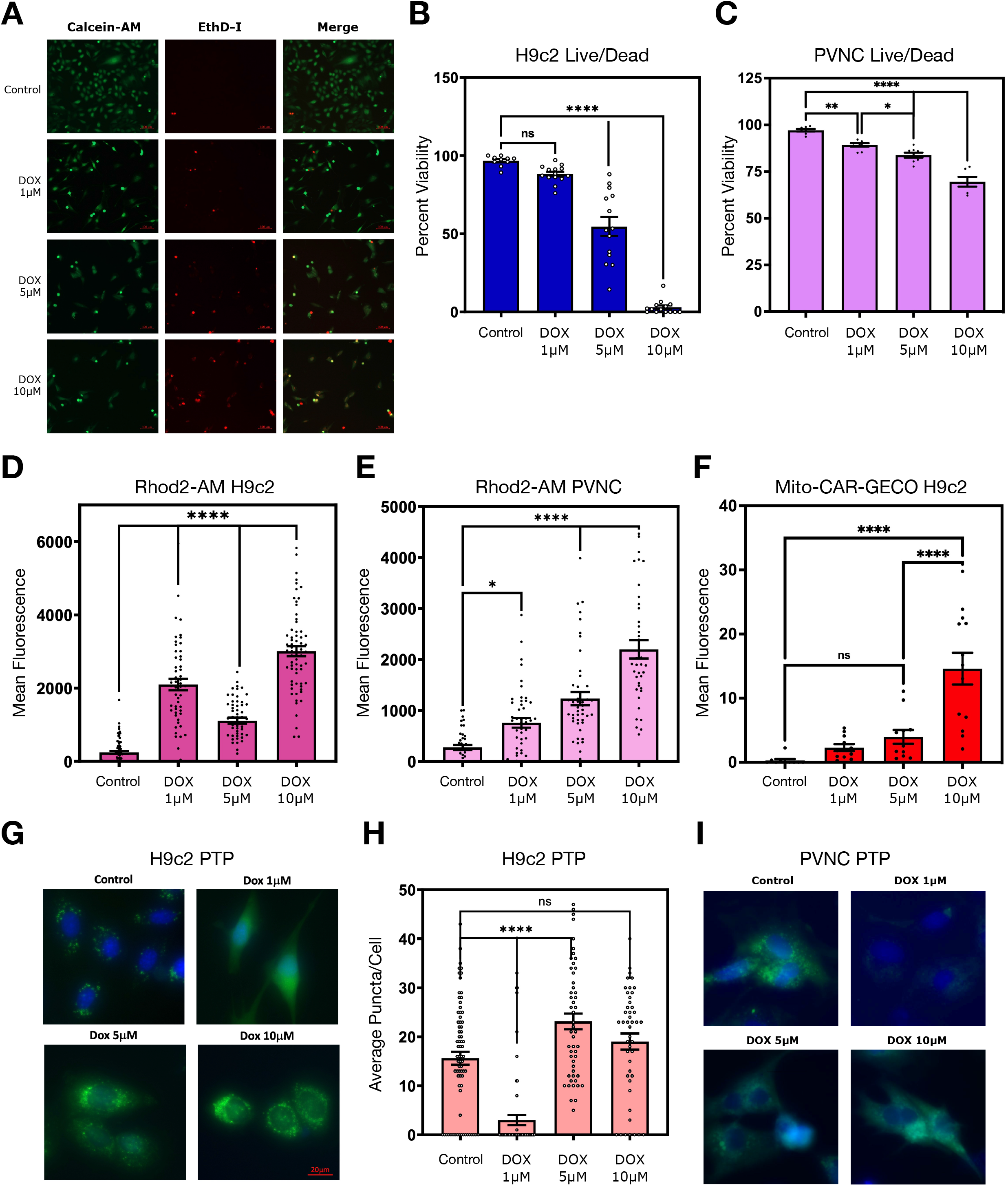
DOX Differentially Impacts Mitochondrial Permeability Transition Despite Elevations in Mitochondrial Calcium. A) H9c2 cells were treated with 1, 5, and 10 μM DOX for 24-hours and stained with calcein-AM (green) and ethidium homodimer-1 (red; EthD-I)) to identify living and dead cells, respectively (Live/dead assay). B) Quantification of the data in (a). C) Quantification of live/dead assay performed in PVNC’s, as described in (a). D) Mean fluorescence quantification of H9c2 cells treated as in (a) and stained with reduced Rhod2-AM (dihydrorhod2). E) Mean fluorescence quantification of PVCN’s treated as in (a) and stained with reduced Rhod2-AM (dihydrorhod2). F) Mean fluorescence quantification of H9c2 cells transfected with the mitochondrial-targeted calcium biosensor, Mito-CAR-GECO, and treated as in (a). G) Representative images of H9c2 cells treated as in (a) and stained with Calcein-AM and cobalt chloride to evaluate mitochondrial permeability transition. H) Quantification of the data in (g) by calculated the average puncta per cell. I) Representative images of PVCN’s treated as in (a) and stained with Calcein-AM and cobalt chloride. All data are represented as mean ± S.E.M. * indicates p<0.05, **p<0.01, ****p<0.0001. ns = not significant.

### DOX Induces Mitochondrial Oxidative Stress, Lipid Peroxidation, and AIF Translocation

Next, we evaluated mitochondrial membrane potential (ΔͰm) by TMRM staining. Similar to the permeability transition studies, we observed that 1 μM DOX reduced ΔͰm, but higher concentrations of DOX had no effect (Figure 5A-B). Next, we evaluated cellular oxidative stress using DCF staining. Shown in Figure 5C, we observed a dose-dependent increase in DCF staining with DOX exposure. Using MitoSOX, we observed that DOX produced a dose-dependent increase in mitochondrial superoxide in H9c2 cells and PVNCs (Figure 5D-F). As mitochondrial permeability transition was not impacted by high concentrations of DOX, but both calcium and reactive oxygen species were increased, we evaluated the subcellular localization of Apoptosis-inducing factor (AIF), a mitochondrial redox sensor implicated in caspase-independent cell death and DNA degradation that has been previously shown to be activated by oxidative stress, accumulation of mitochondrial calcium and Calpain cleavage, and PARP1 (38–41). In control PVNCs, we observed a mitochondrial distribution of AIF. However, in DOX treated cells we observed a dose-dependent nuclear accumulation of AIF with concurrent dissipation of the nuclear stain (Figure 5G-H). We also evaluated mitochondrial lipid peroxidation using the MitoPeDPP stain. DOX-induced increase in mitochondrial lipid peroxidation that was particularly apparent at 10 μM (Figure 5I). Finally, we evaluated the subcellular locational of HMGB1, a nuclear protein that is released into the cytosol and extracellular space during necrosis where it acts as an alarmin or damage-associated molecular pattern (DAMP) to engage an inflammatory response. Intriguingly, DOX treatment increased nuclear HMGB1 staining without observable cytosolic staining, suggesting that some aspects of the typical necrotic phenotype are inhibited by DOX at high concentrations (Figure 5J-K).

**Figure 5.**
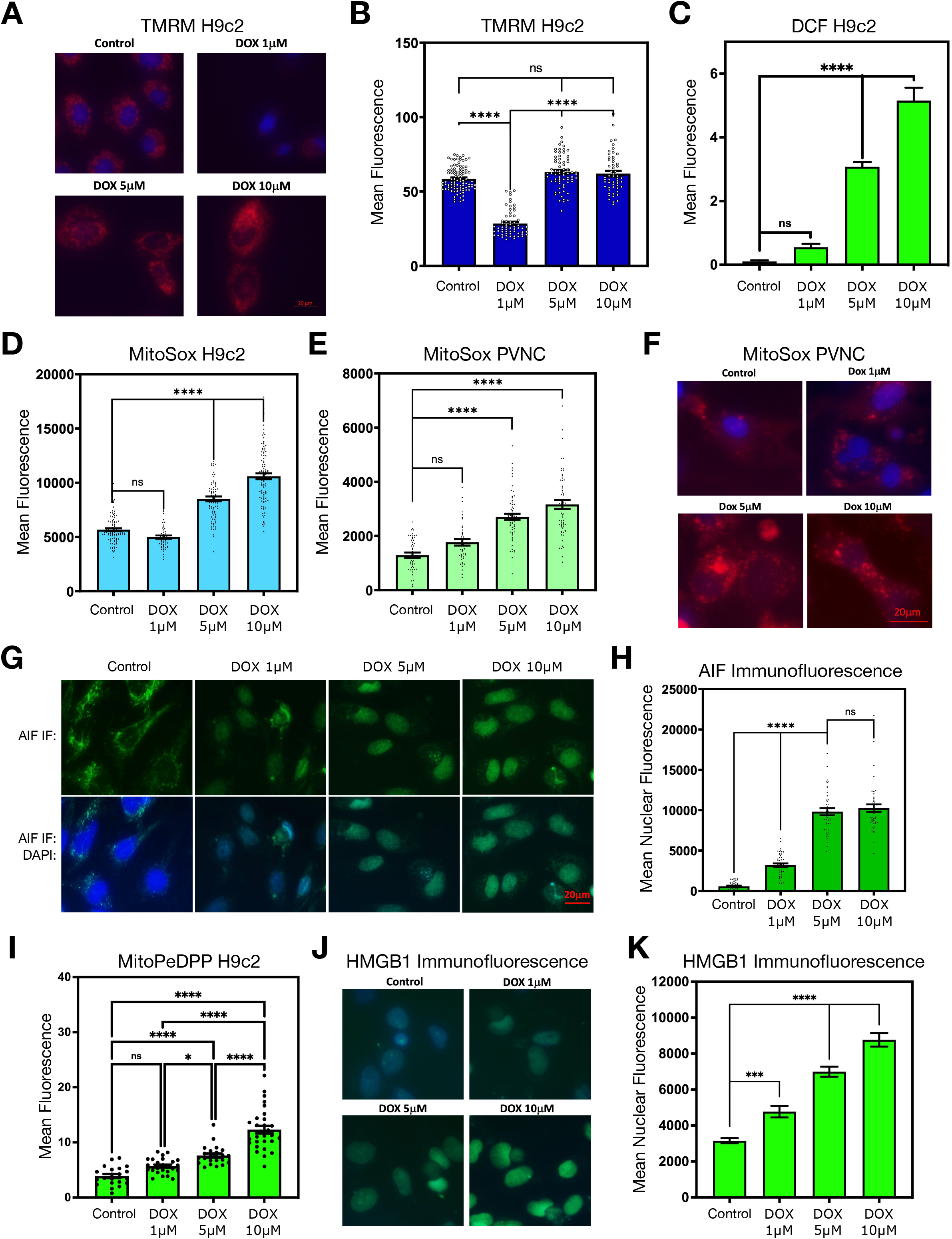
DOX Induces Mitochondrial Oxidative Stress, Lipid Peroxidation, and AIF Translocation. A) Representative images of H9c2 cells were treated with 1, 5, and 10 μM DOX for 24-hours and stained with TMRM to evaluate mitochondrial membrane potential. B) Quantification of the data presented in (a). C) Mean fluorescent quantification of H9c2 cells treated as in (a) and stained with H_2_-DCFDA (DCF) as an indicator for reactive oxygen species accumulation. D) Mean fluorescent quantification of H9c2 cells treated as in (a) and stained with Mito-Sox as an indicator of mitochondrial superoxide. E) Mean fluorescent quantification of PVNC’s cells treated as in (a) and stained with Mito-Sox. F) Representative images of the data presented in (e). G) Immunofluorescence of AIF in PVNC’s, treated as in (a), and counter stained with DAPI. H) Quantification of mean nuclear fluorescence of the data presented in (g). I) Quantification of H9c2 cells treated as in (a) and stained with MitoPeDPP as an indicator of mitochondrial monitor lipid peroxidation. J) Immunofluorescence of HMGB1 in PVNC’s treated, as in (a), and counter stained with DAPI. K) Quantification of mean nuclear fluorescence of the data presented in (j). All data are represented as mean ± S.E.M. * indicates p<0.05, ***p<0.001, ****p<0.0001. ns = not significant.

### DOX Inhibits Autophagic Flux and Activates Mitophagy

Previous studies have implicated autophagy as a necessary pathway to release HMGB1 from the nucleus during necrosis and ferroptosis (42). Thus, we evaluated markers of autophagy and mitophagy in the hearts of DOX treated mice. In mice treated with LDD, markers of autophagy such as LC3 and p62 were unchanged compared to control (Figure 6A). However, in HDD treated mice we observed a marked increase in both LC3-II and p62. As p62 protein levels typically decrease by lysosomal degradation when autophagic flux is increased, we interpreted these data to suggest that autophagic flux was inhibited by HDD. This observation may explain why HMGB1 accumulated in the nucleus of PVNCs exposed to high concentrations of DOX. We also observed a modest increase in Parkin and BCL-2 expression, but these did not achieve significance. Next, we performed dose-response experiments in H9c2 cells. Shown in Figure 6C-D, we observed a dose-dependent increase in LysoTracker staining indicating that lysosomal acidification was not impaired at higher concentrations of DOX. In addition, we monitored LC3-GFP and observed autophagosome formation at every dose of DOX; however, at higher concentrations of DOX there was increased co-localization of LC3-GFP with LysoTracker staining (Figure 6C). As BNIP3 and Parkin are known regulators of mitochondrial autophagy (ie. Mitophagy)(43–45), we utilized the mito-pHred mitophagy biosensor in DOX exposed cells. DOX exposure produced increased levels of mitophagy with increasing dose (Figure 6E-F). Finally, we evaluated the subcellular localization of p53, as this tumour suppressor gene has been previously shown to respond to genotoxic stress, is a dual regulator of autophagy, and can elicit both apoptotic and necrotic cell death (46–49). Shown in Figure 6G-H, DOX induced the expression of p53 which accumulated in the nucleus.

**Figure 6.**
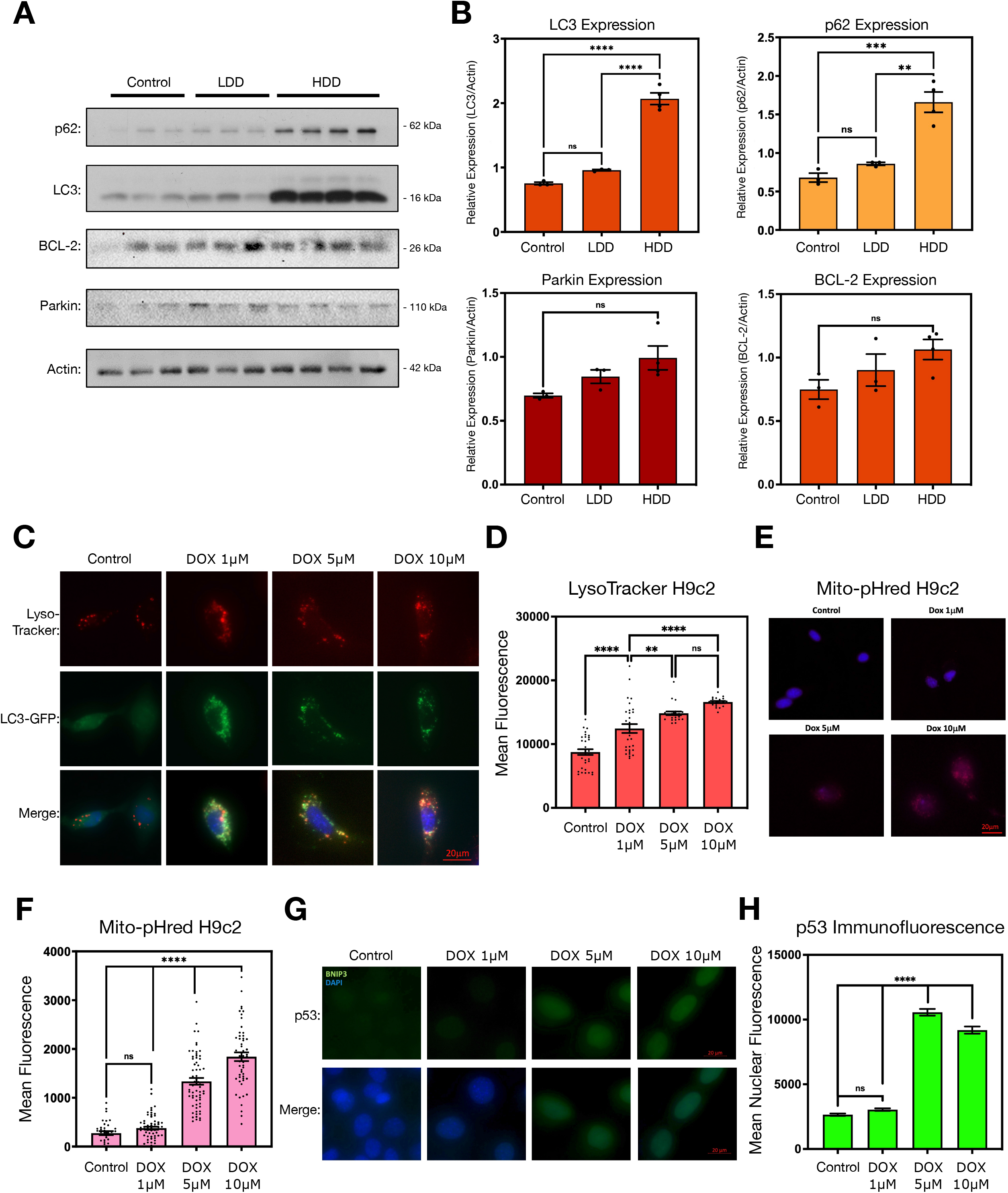
DOX Inhibits Autophagic Flux and Activates Mitophagy. A) Western blot analysis of control, low-dose DOX (LDD; 8 mg/kg), and high-dose DOX (HDD; 20 mg/kg) treated mice at PID3, including p62, LC3, BCL-2, Parkin, and β-actin (Actin). B) Quantification of western blots in (a) relative to β-actin (Actin). C) Representative images of H9c2 cells transfected with LC3-GFP plasmid (green; LC3-GFP), then treated with 1, 5, and 10 μM DOX for 24-hours and stained with LysoTracker (red; Lyso-Tracker) to monitor autophagolysosome formation, and counterstained with Hoechst (blue). D) Mean florescent quantification of LysoTracker data presented in (c). E) Representative images of H9c2 cells transfected with Mito-pHred and then treated with DOX as in (c). F) Mean fluorescent quantification of data presented in (e). Immunofluorescence of p53 in H9c2 cells treated with DOX as in (c) and counterstained with DAPI. H) Mean fluorescent quantification of data presented in (g). All data are represented as mean ± S.E.M. * indicates p<0.05, ***p<0.001, ****p<0.0001. ns = not significant.

### Pharmacological Blockade of Autophagic Flux Promotes a Necrotic Phenotype in DOX Exposed Cells

Next, we sought to determine the cellular consequences of impaired autophagic flux and the inhibition of cell death pathways on DOX-exposed H9c2 cells. Using Annexin-V (Ann) and propidium iodide (PI) staining, we performed flow cytometry analysis to identify viable (Ann-/PI-), apoptotic (Ann+), necrotic/late apoptotic (Ann+/PI+), and necrotic (PI+) cells. We observed that DOX exposure produced a dose-dependent decline in cell viability (Figure 7A). In addition, we observed an increase in apoptotic cells (Ann+) at 1 and 5 μM DOX; however, at 1 μM DOX nearly 30% of cells were PI+, and at 5 μM DOX approximately 70% of cells were PI+ (Figure 7A). To pharmacologically inhibit autophagic flux, we treated cells with Bafalomycin-A1 (Baf) and evaluated the impact on cell viability when cells were exposed to DOX (Figure 7B). Baf treatment reduced cell viability at every dose of DOX, but not in control cells. In addition, Baf treatment increased the percentage of Ann+/PI+ cells, suggesting that inhibition of autophagy increased cell death where plasma membrane integrity was compromised (Figure 7B).

**Figure 7.**
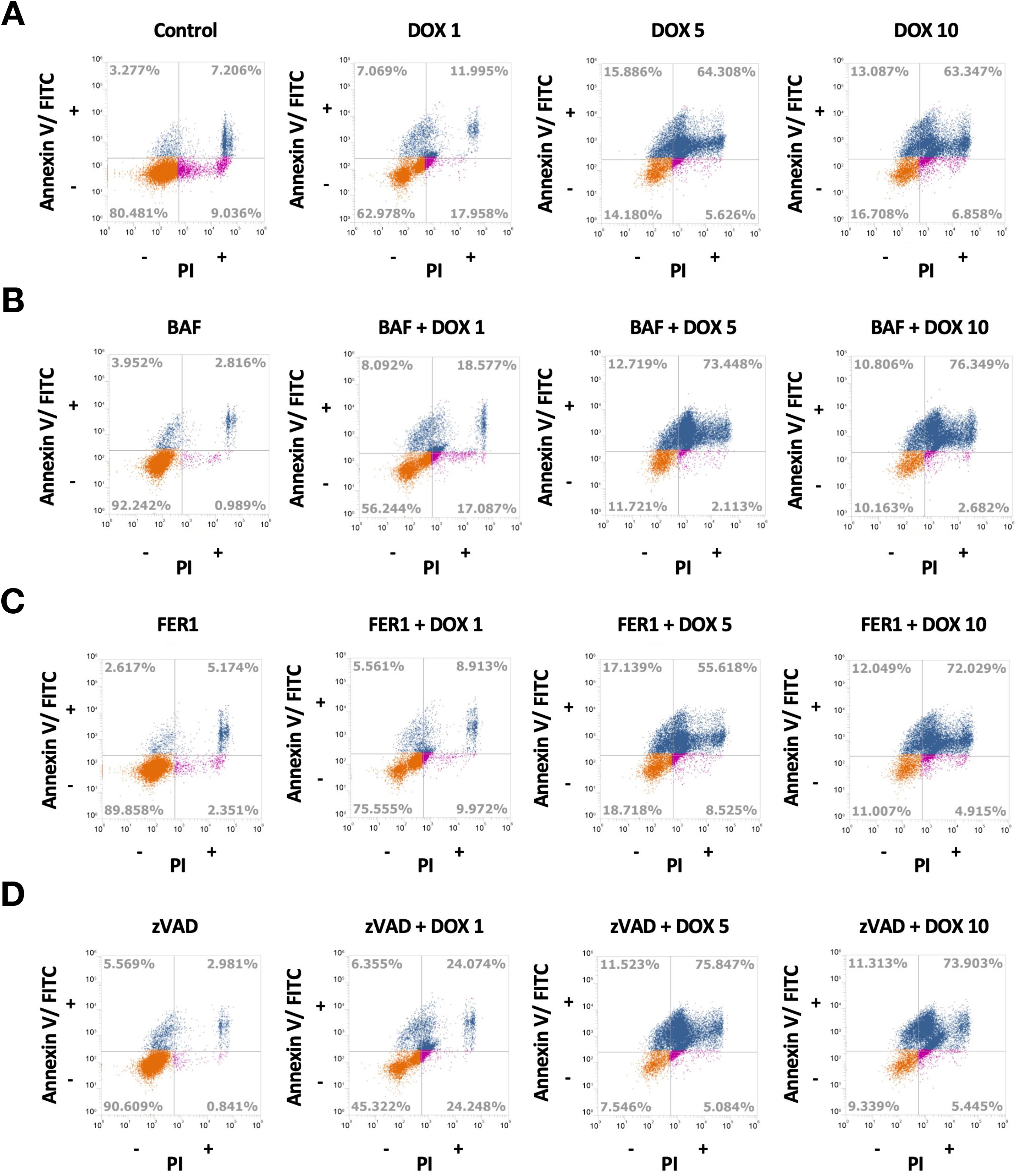
Pharmacological Blockade of Autophagic Flux Promotes a Necrotic Phenotype in DOX Exposed Cells. A) Representative dot-plots of H9c2 cells that were treated with 1, 5, and 10 μM DOX for 24-hours and then incubated with propidium iodide (PI) and Annexin V (Annexin V/FITC) before analysis with a flow cytometer. B) Representative dot-plots of H9c2 cells that were pre-treated with 5nM Bafilomycin A1 (BAF) and DOX as in (a). C) Representative dot-plots of H9c2 cells that were pre-treated with 1μM Ferrostatin-1 (FER1) and DOX as in (a). D) Representative dot-plots of H9c2 cells that were pre-treated with 50μM zVAD(OMe)-fmk (ZVAD) and DOX as in (a). Each condition was performed in triplicates and recorded 20,000 events.

As we observed activation of caspase-3 at low concentrations of DOX, and increased levels of mitochondrial lipid peroxidation at higher concentrations of DOX, we combined DOX exposure with zVAD-fmk (zVAD), a pan-caspase inhibitor, and Ferostatin-1 (FER1) an inhibitor of ferroptosis. When combined with DOX exposure, FER1 decreased the percentage of Ann+/PI+ cells at 1 μM and 5 μM DOX and improved cell viability at these DOX concentrations (Figure 7C). Intriguingly, zVAD treatment decreased the percentage of Ann+ cells at every dose of DOX, but did not improve cell viability (Figure 7D), suggesting that when apoptotic pathways were inhibited, cell death was achieved through alternative necrotic pathways. Collectively, these observations suggest that inhibiting autophagy flux in DOX-exposed cells increases necrotic cell death; whereas, inhibiting lipid peroxidation with FER1 produces some improvement in cell viability up to concentrations of 5 μM DOX.

## Discussion

DOX-induced cardiotoxicity can present acute, early, or late (50); however, patients experiencing acute-onset cardiotoxicity usually recover within a week (51). These observations in human trials are in stark contrast to rodent studies where HDD regimes produce a 50-60% mortality within 5-days post-exposure (19, 20). In the present study, we undertook detailed structure/function and cellular analysis of acute LDD and HDD exposure using 3 independent model systems. Our observations suggest that acute HDD elicits substantial cardiac necrosis, diminished cardiac output with reduced left ventricular diastolic volume, combined with inhibition of autophagic flux and increased BNIP3 expression. These findings suggest that acute HDD does not recapitulate the disease phenotype observed in acute human cardiotoxicity, which often presents with an increase in the left ventricular end-diastolic diameter, nor does acute HDD exposure generate the characteristic alterations of late DOX-induced cardiotoxicity with respect to cardiac fibrosis or systolic dysfunction.

Although recent literature has identified several novel forms of necrotic cell death in the heart (15), and early guidelines have been developed to promote consistent evaluation and interpretation of these forms of cell death (52), the observations in the present study suggest that necroptosis, pyroptosis, and ferroptosis may only play a minor role during acute HDD cardiotoxicity. However, our results are consistent with the work of others regarding the role of autophagy in DOX-induced cardiotoxicity (11, 22, 23). Autophagic cell death can occur through either excessive autophagic flux or a blockade of autophagic flux (46, 53, 54). Our observations support the notion that impaired autophagic flux contributes to DOX-induced necrosis; however, our data from cell culture suggests that lysosomal acidification (determined through lysotracker staining) and acidification of mitochondrial fragments (determined by mito-pHred fluorescence) continues even at the highest tested DOX concentration. Although not conclusive, these data suggest that the inhibition of autophagic flux may involve multiple mechanisms beyond impaired lysosomal acidification (22). Previous studies have shown that nuclear p53 typically drives the expression of apoptotic genes and genes that activate autophagy, compare to cytoplasmic/mitochondrial p53 which can inhibit autophagy and trigger necrosis (46). Our culture data identifies that p53 accumulates in the nucleus following DOX exposure, suggesting that p53 expression may not be responsible for impaired autophagy observed by HDD. Our results are also consistent with the work of other laboratories demonstrating that BNIP3 expression is increased with acute HDD exposure (19). However, BNIP3 expression was not altered in the more dose appropriate chronic DOX model suggesting that not all changes in gene expression are conserved between acute HDD and cumulative DOX exposure models.

Human studies have the determined that the most significant risk factor predicting DOX-induced cardiotoxicity is cumulative dose (50). However, based on our observations, and the work of others, it appears that this premise has been taken out of context and incorrectly extrapolated to imply that exposure to a large dose of DOX acutely will result in the same cardiac phenotype as repeated smaller doses of DOX over a longer period of time, consistent with the human chemotherapeutic regimen. Notwithstanding, chemotherapy doses are typically higher in pediatric oncology, and the acute HDD model may have some validity as an early life model of DOX exposure with the goal of evaluating cardiac function later in life. In this regard, treatment of a childhood cancer with DOX is one of the most pronounced risk factors for adult-onset cardiotoxicity and heart failure (50). However, animal studies have shown that this early life DOX exposure phenomenon can be recapitulated by exposing mice to 1 mg/kg at postnatal day 5 and 10, and then 0.5 mg/kg at postnatal day 15 and 20 (55). Using this protocol, researchers identified cardiotoxicity by 12-weeks of age, including impaired exercise tolerance and cardiac maladaptation, and a worsening prognosis and increased infarct area following coronary ligation. Mechanistically, this phenotype was attributed to fewer cardiac progenitor cells following DOX exposure, and impaired cardiac blood vessel formation and regenerative capacity. These findings suggest that a cumulative dose of 3 mg/kg over the course of 15 days in early life is sufficient to approximate the pediatric DOX phenotype, and also suggest that an HDD routine involving 20 mg/mL is beyond pharmacological.

There are a number of limitations to the present study that will require further investigation. Firstly, we chose to use male mice as most animal studies evaluating DOX-induced cardiotoxicity use this sex. Additional work is needed to determine if this acute phenotype is consistent in female mice, or if biological sex differences exist. Secondly, our sample size was only large enough to statistically analyze effects of substantial magnitude, and a larger study would be required to evaluate more nuanced effects. Thirdly, our study was limited to C57BL/6 mice of the -N sub-strain. As a number of metabolic differences have been reported between the N an J sub-strains, addition investigation is needed to determine if the reported phenotype is conserved among sub-strains and in other rodent models. Finally, additional work is required to evaluate DOX-induced cardiotoxicity models and carefully compare existing models to human cardiotoxicity, as has been recently done for ischemic heart disease models (56).

In summary, we describe the effects of acute LDD and HDD DOX on cardiac structure/function, cell death and autophagy pathways. Echocardiographic analysis revealed that HDD acutely causes a diminished cardiac output and left ventricular volume, without impacting ejection fraction or fractional shortening. In addition, LDD activated caspase signalling, while acute HDD inhibited autophagic flux and increased BNIP3 expression. Collectively, these observations help characterize the structure/function and gene expression alterations in the mouse heart during the acute phase of DOX-induced injury, but also suggest that acute HDD does not recapitulate the disease phenotype of human cardiotoxicity and that not all changes in gene expression are conserved between acute exposure and chronic cumulative DOX models.

## Acknowledgements

This work was support by a Discovery Grant from the Natural Science and Engineering Research Council (NSERC) Canada, a Grant-in-Aid from the Heart and Stroke Foundation of Canada, and an operating grant from the Children’s Hospital Research Institute of Manitoba to JWG. VWD is supported by a Grant-in-Aid from the Heart and Stroke Foundation of Canada and is the Allen Rouse Basic Scientist of the Manitoba Medical Services Foundation. SG is supported by the Health Sciences Foundation, Winnipeg, Canada, and an operating grant from the Children’s Hospital Research Institute of Manitoba. PK was supported through an NSERC Summer studentship and a stipend from the University of Manitoba Bachelor of Science in Medicine program. MDM and WM were supported by studentships from the Children’s Hospital Foundation of Manitoba and Research Manitoba. JTF was supported by an Alexander Graham Bell studentship from NSERC Canada. AMC was supported by a University of Manitoba Undergraduate Research Award.

